# Grooming as a window into the post-stress recuperative process

**DOI:** 10.64898/2026.05.27.728357

**Authors:** Afra N Mahmud, Ann K PierreLouis, Naomi Yamaguchi, Denise J Cai, Zachary T Pennington

## Abstract

Alterations in rodent self-grooming have been used to model various facets of neuropsychiatric illness. In the context of affective behavior, increases in grooming have been proposed as a sign of stress. This is because grooming has been observed to increase in close temporal proximity to stressful events. However, in other situations, stress appears to suppress grooming, complicating the utility of measuring grooming in the study of stress and mental health. Here, we show that this discrepancy can be resolved by considering time and experimental context. We found that in initial response to stress, grooming declined in proportion to stressor intensity. Moreover, stress-related cues and anxiogenic stimuli similarly suppressed grooming. Conversely, optogenetic inhibition of the amygdala in a stress-associated context decreased threat-elicited freezing, consistent with a reduction in stress, and increased grooming. These results indicate that the immediate response to stress is a suppression of grooming. However, when stressed mice were returned to their homecage environment, grooming increased. Similarly, mice increased grooming when they returned to the safe zone in an anxiety assay. Accordingly, rather than being a defensive response to signs of danger, increased grooming seems to reflect a post-stress response that occurs once animals detect the absence of danger. These findings suggest that post-stress grooming could provide a window into the neurobiology of post-stress recuperative processes.

## INTRODUCTION

Grooming is an evolutionarily conserved behavior that supports a range of vital functions, from guarding individuals from pathogens to influencing social interactions [1]. Importantly, alterations in rodent self-grooming behavior have been proposed to reflect various processes relevant to studying the neurobiology of mental health [2]. For example, increased self-directed grooming in genetic autism models has been postulated to capture repetitive behavioral symptoms [2–5], and increased repetitive grooming has been used to model stereotyped behaviors in obsessive compulsive disorder [6].

Grooming has also been closely tied to stress and affective behavior, with several proposing that grooming might serve as a measure of an animal’s affective or stress state [2,4,7]. This is because grooming has been observed to increase following the presentation of a range of acute stressors, including restraint [8,9], footshock [10], testing in anxiety-related behavior assays such as the elevated plus maze [11], exposure to novel environments [12,13], exposure to bright lights [14,15], and exposure to social stressors [11,12]. Additionally, animals that have experienced chronic social defeat stress in the past display elevated grooming [16], as do some rodent strains with high trait anxiety [17,18] (although discrepancies also exist [13,19]). These findings broadly suggest that heightened grooming might be useful to infer stress levels.

Although much evidence indicates that stress promotes grooming in rodents, several findings suggest otherwise. For example, novel environments [20], footshock [10,19], cat urine [21], as well as cats themselves [22] have been shown to suppress grooming levels. These findings indicate that grooming cannot be used as a simple readout of stress, but rather, must be considered in the context of environmental and/or individual modulators. Unfortunately, what these factors are remain unclear. Critically, this knowledge gap renders measures of grooming difficult to interpret.

Here, we examined whether time and experimental context may allow these discrepancies to be resolved. Most, if not all, studies that have found that stress decreases grooming have observed these changes in the presence of the stressor [10,19–22]. Conversely, studies that have found stress increases grooming have measured grooming after stress, and in a context distinct from where the stressor occurred [8,9,11,14,15,23]. This led us to hypothesize that grooming serves as a post-stress behavior, possibly used to restore animals to a pre-stress state. Using a range of stressor types and stressor levels, and recording animals both during and after stress, we systematically tested this hypothesis. Our results demonstrate that the immediate response to stress is a decrease in grooming, whereas grooming increases when mice are removed from a stressful environment. These findings are readily integrated with existing ethological accounts of defensive behavior and can support improved understanding of fluctuations in grooming when modeling aspects of neuropsychiatric function.

## METHODS

### Animals

Animals were adult C57BL/6J mice obtained from Jackson Laboratories, aged 2-6 months. An equal mix of male and female mice was used for all experiments, except for the experiment testing the impact of optogenetically inhibiting the amygdala, for which only male mice were used. Group sizes and the number of mice per sex are listed in each figure legend. Mice were singly housed in a temperature- and humidity-controlled vivarium on a 12/12 light-dark cycle (lights on at 7 a.m.), and all handling and behavioral testing took place during the light phase. All experimental procedures were approved by the Icahn School of Medicine at Mount Sinai’s IACUC.

### Surgery

For surgery, anesthesia was induced with 5% isoflurane and subsequently maintained at 1-2%. Body temperature was maintained both during surgery and surgical recovery with a heating pad below the mice, and ophthalmic ointment was applied to lubricate the eyes. 150 nL of virus was infused bilaterally into the basolateral amygdala (BLA) at the following coordinates relative to bregma: AP: −1.4; ML: 3.3; DV: −5 (mm). Virus was infused via glass micropipettes at 2 nL/second and pipettes were left in place for 5 minutes before retraction. Mice received a viral cocktail with AAV9-hSyn-Cre-hGH (final titer: 1 × 10^12^ GC/mL; Addgene #105555) and AAV5-hSyn-SIO-stGtACR1-FusionRed (final titer: 1.89 × 10^12^ GC/mL; Addgene #105678), or AAV9-hSyn-Cre-hGH (final titer: 1 × 10^12^ GC/mL; Addgene #105555) and AAV5-hSyn-DIO-mCherry (final titer: 1.98 × 10^12^ GC/mL; Addgene #50459). Fiberoptic cannulas (200 *μ*m, 0.5 NA, RWD Life Sciences, #807-00046-00) were lowered to 0.4 mm above the infusion site and affixed to the skull with super glue and black dental cement. Following surgery, mice were given 20 mg/kg ampicillin and 5 mg/kg carprofen (s.c.) daily for 7 days, and body weight and general disposition were monitored.

### Behavior

#### Habituation

Prior to all behavioral experiments, mice were habituated to gentle handling for 1 min/day over 5-7 days. Additionally, on 3 of these days, mice were transported to the experimental testing area and handled there. For optogenetic experiments, mice were additionally habituated to being connected to fiberoptic patch cords for 3 days, for approximately 2 minutes per day.

#### Footshock conditioning/recall

During conditioning, mice were placed in a conditioning chamber housed within a sound-attenuating cubicle (Med Associates), and after a 5-minute exploration period, received a series of 10 footshocks (0.25 mA or 1 mA). Each shock was 1 second in duration and was separated by 30 seconds. Mice were taken out of the chamber 30 seconds after the last shock and returned to their homecage. In total, each session was 10 minutes and 10 seconds in length. For recall sessions, mice were returned to the conditioning chamber for 8 minutes. Non-shocked mice underwent the same procedures but did not receive footshocks. Mice were recorded by an infrared camera mounted to the inside of the sound-attenuating chamber.

#### Post-stress recordings

Post-stress recordings took place in the mice’ homecage immediately after the footshock conditioning procedure described above. The homecage was set on a rack in the conditioning room but was separated from the rest of the testing room by a heavy black curtain hung from the ceiling. Each cage was recorded using two cameras positioned approximately 8 inches from two sides of the cage (providing side and front views). Each recording was 10 minutes in length and began immediately following conditioning. Prior to conditioning, mice were habituated to sitting in their cages on the rack for approximately 30 minutes per day for 3 days. Lighting on the rack was 40-50 lux.

### Light-Dark/Dark-Dark testing

All testing took place in an acrylic chamber (30.5 × 24.1 × 21 cm) bisected by a wall containing a small opening to connect the chamber’s 2 sides (A and B). The top of the chamber was transparent to permit illumination via a combination of white and infrared lights above the chamber, and the front of the chamber was transparent to allow recording via an infrared camera located just outside. The chamber walls were otherwise opaque. Sides A and B were equivalent except for their flooring: one side had a smooth plastic floor; the other had a plastic floor with small horizontal grooves. Sides A and B could be independently covered with red transparency film to make them light or dark (from 400 lumens uncovered to 10 lumens covered). Side B was always dark.

Testing took place across 3 days: habituation, Light-Dark, and Dark-Dark. The order of Light-Dark and Dark-Dark testing was counterbalanced across mice. During habituation, mice were placed on Side B (dark) for 20 minutes, with a barrier preventing the mice from entering or seeing Side A. This session served to make Side B familiar to the mouse. Then, during Light-Dark and Dark-Dark testing, mice were initially placed in Side B but allowed to roam freely between sides A and B for 8 minutes. Side A was well lit in the Light-Dark test and dark during the Dark-Dark test.

#### Optogenetics

For stGtACR1-mediated inhibition, laser light was delivered with a 473 nM laser (OptoEngine LLC) connected via a patch cord to a bifurcating rotary joint (Doric). Consistent with prior reports utilizing stGtACR1 in the amygdala and hypothalamus [24,25], we utilized 20 Hz, 20 ms pulse width, 5 mW, illumination. Light intensity was measured from the fiber tip using a light meter (PM100D with S130C attachment, ThorLabs). During recall tests, light was delivered in an OFF-ON-OFF-ON pattern (2 minutes each). The average time freezing when the light was ON and OFF is presented.

### Quantification

#### Grooming

All videos were scored post-hoc by observers that were generally blind to experimental condition (except when mice were being shocked in the video). For scoring purposes, videos were binned into 2 second increments. For each bin, observers scored whether any grooming occurred. Accordingly, the proportion of bins where grooming occurred is presented. The following events were classified as grooming: circular swiping of head/face with the forepaws and/or hindpaws, grooming of the flank/hind quarters with the mouth, or grooming of the tail. Inter-rater reliability was assessed for a subset of videos and the average correlation between 3 raters was found to be above 0.95.

#### Freezing/Position

Freezing of untethered mice was scored using Med Associates VideoFreeze Software [26]. For mice tethered by optogenetic patch cords, ezTrack was used [27]. The position of mice within the Light-Dark and Dark-Dark tests was scored using ezTrack [28].

### Histology

Following behavioral testing, mice that underwent surgical manipulation were deeply anesthetized, transcardially perfused with 5 mL 1X PBS followed by 5 mL 4% PFA, and their brains were then extracted and placed in paraformaldehyde at 4C overnight. The next day, brains were transferred to 30% sucrose in 1X PBS, left at 4C to sink before being frozen, and then sectioned at 50 μm on a cryostat. Lastly, tissue was mounted onto slides, cover-slipped using mounting media with DAPI (Vector Laboratories, #H-1200-10), and imaged on a Leica DM6 epifluorescent microscope. Viral expression and cannula placement were evaluated using the mouse brain atlas of Franklin and Paxinos [29].

### Statistical analysis

All analyses were performed using RStudio. Briefly, omnibus ANOVA were conducted using the ezANOVA package with Type III degrees of freedom. The white adjustment was implemented to correct for heterogeneity of variance using heteroscedasticity corrected standard errors (‘hc3’). For repeated measures ANOVA, the Greenhouse-Geisser correction was implemented when the assumption of sphericity was not met. Simple effects were only examined following significant omnibus interactions, but omnibus tests are not reported in the main text for the sake of clarity. Detailed statistics with omnibus results can be found in Supplementary Table 1. F-values and t-values are rounded to the nearest tenth and hundredth, respectively. Where F values were less than .1, F is listed as 0. All figures depict the mean and standard error of the mean. For all figures, * = p<0.05, ** = p<=.01, *** = p<=0.001.

## RESULTS

### Grooming is immediately suppressed by stressors and stress-associated cues

We first sought to assess the relationship between acute stress and grooming, parametrically varying stressor intensity. This allowed us to begin to evaluate whether different levels of stress might uniquely influence grooming – for example, if low levels of stress increase grooming and high levels of stress decrease grooming, as had been previously suggested [30]. Mice underwent conditioning in which they received varying intensities of electric footshock (Fig 1A). After a five-minute baseline/preshock exploration period in an experimental testing chamber, mice received either No Stress (no footshock), Low Stress (10, 0.25 mA, footshocks), or High Stress (10, 1 mA, footshocks). As expected, Low Stress and High Stress mice displayed increased levels of defensive freezing over the course of the session (Fig 1B). Additionally, mice in the High Stress group froze more than mice in the Low Stress group, demonstrating that low and high intensity footshocks indeed elicited varying levels of stress (Fig 1B). Next, examining grooming during this session, the 3 groups did not show differences in grooming during the preshock period (Fig 1C). However, in the 5 minutes after shock commenced, both Low Stress and High Stress mice showed a near complete suppression of grooming (Fig 1C). By contrast, the grooming of No Stress mice did not change from the preshock period to the shock period.

**Figure 1.**
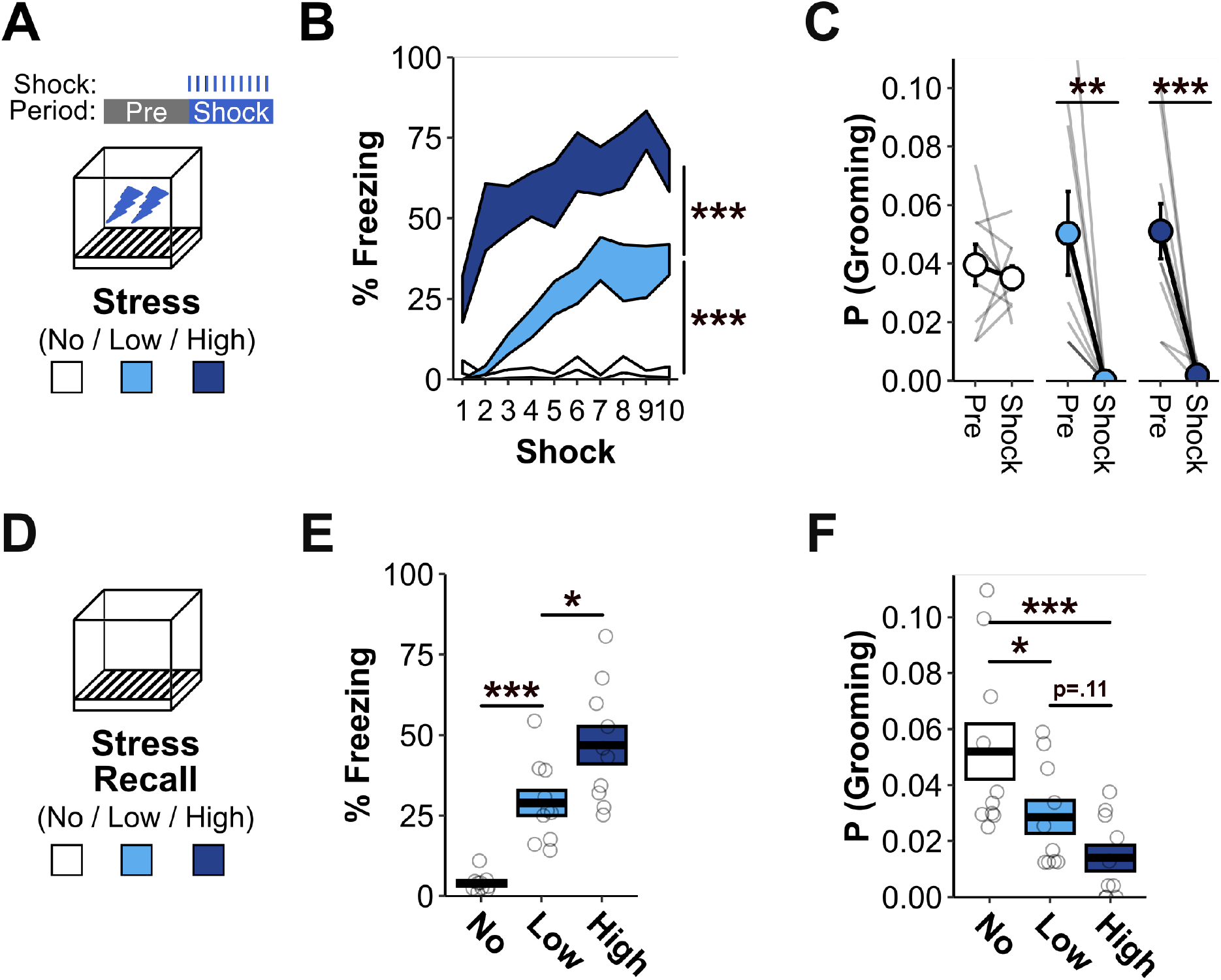
Stress and stress-associated cues suppress grooming. A) Mice were exposed to No, Low, or High stress (no shocks, 10 weak shocks, or 10 strong shocks). No Stress: N=10 (5 females); Low Stress: N=10 (5 females); High Stress: N=10 (5 females). B) Post-shock freezing increased in stressed mice and differed between Low Stress and High Stress groups. No Stress vs Low Stress: F_1,16_=39.1, p<0.001; No Stress vs High Stress: F_1,16_=106.9, p<0.001. Low Stress vs High Stress: F_1,16_=32.8, p<0.001. C) Grooming was suppressed by both Low Stress and High Stress. Effect of Time for each group: No Stress: F_1,7_=0.3, p=0.63; Low Stress: F_1,8_=12.7, p<0.01; High Stress: F_1,8_=23.9, p=0.001). Effect of Group on preshock/baseline grooming: F_2,23_=0.4, p=0.65. D) The same mice from A-C were returned to the stressor environment for a recall session. E) During recall, freezing increased in proportion with stress severity. No Stress vs Low Stress: F_1,16_=29.3, p<0.001; No Stress vs High Stress: F_1,16_=39.5, p<0.001. Low Stress vs High Stress: F_1,16_=4.9, p=0.04. F) During recall, grooming decreased in proportion with stress severity. No Stress vs Low Stress: F_1,16_=5.2, p=0.04; No Stress vs High Stress: F_1,16_=17.9, p<0.001. Low Stress vs High Stress: F_1,16_=3, p=0.11.

The preceding findings indicate that the acute experience of stress in the form of footshock produces a marked suppression of grooming. However, it could be argued that footshock represents a particularly high level of stress. Additionally, the suppression of grooming by footshock may be attributable to shock-elicited escape behaviors interfering with grooming. To address these possibilities, a week later we returned the same mice to the shock-paired environment for a stressor recall session, this time without shock (Fig 1D). This allowed us to assess how a stress-predictive cue that is not physically painful influences grooming. Again, freezing increased in proportion to the intensity of stress previously experienced in the chamber (Fig 1E). Conversely, grooming decreased in proportion to stressor intensity (Fig 1F). Therefore, it appears that both acute stress and stress-predictive environments suppress grooming, and this suppression is proportional to stressor intensity.

### Optogenetic inhibition of stress behaviors restores grooming

The prior findings suggest that increased levels of stress suppress grooming. To test whether this relationship is bidirectional, we assessed if reducing stress levels would increase grooming. To do so, we optogenetically silenced the basolateral amygdala (BLA), a region known to store associative memories for stress-predictive cues. A virus expressing the inhibitory opsin stGtACR1, or a control virus, was infused in the BLA (Fig 2A-B). Optic fibers were then implanted overlying the BLA, such that blue light could be administered to suppress neuronal activity. A month later, mice were conditioned with 10, 1 mA footshocks (Fig 2C). Freezing increased across the course of the session and did not differ between viral groups (Fig 2D). Mice were then returned to this environment 2 days later for a recall test. During the recall test, blue light was alternated OFF and ON to transiently inhibit the BLA. In line with prior reports, optogenetic inhibition of the BLA reduced freezing levels, consistent with a suppression of the stress response (Fig 2E). Conversely, optogenetic inhibition of the BLA produced a stark increase in grooming (Fig 2F). Accordingly, in addition to stress suppressing grooming, neural manipulations that reduce cue-evoked stress increase grooming.

**Figure 2.**
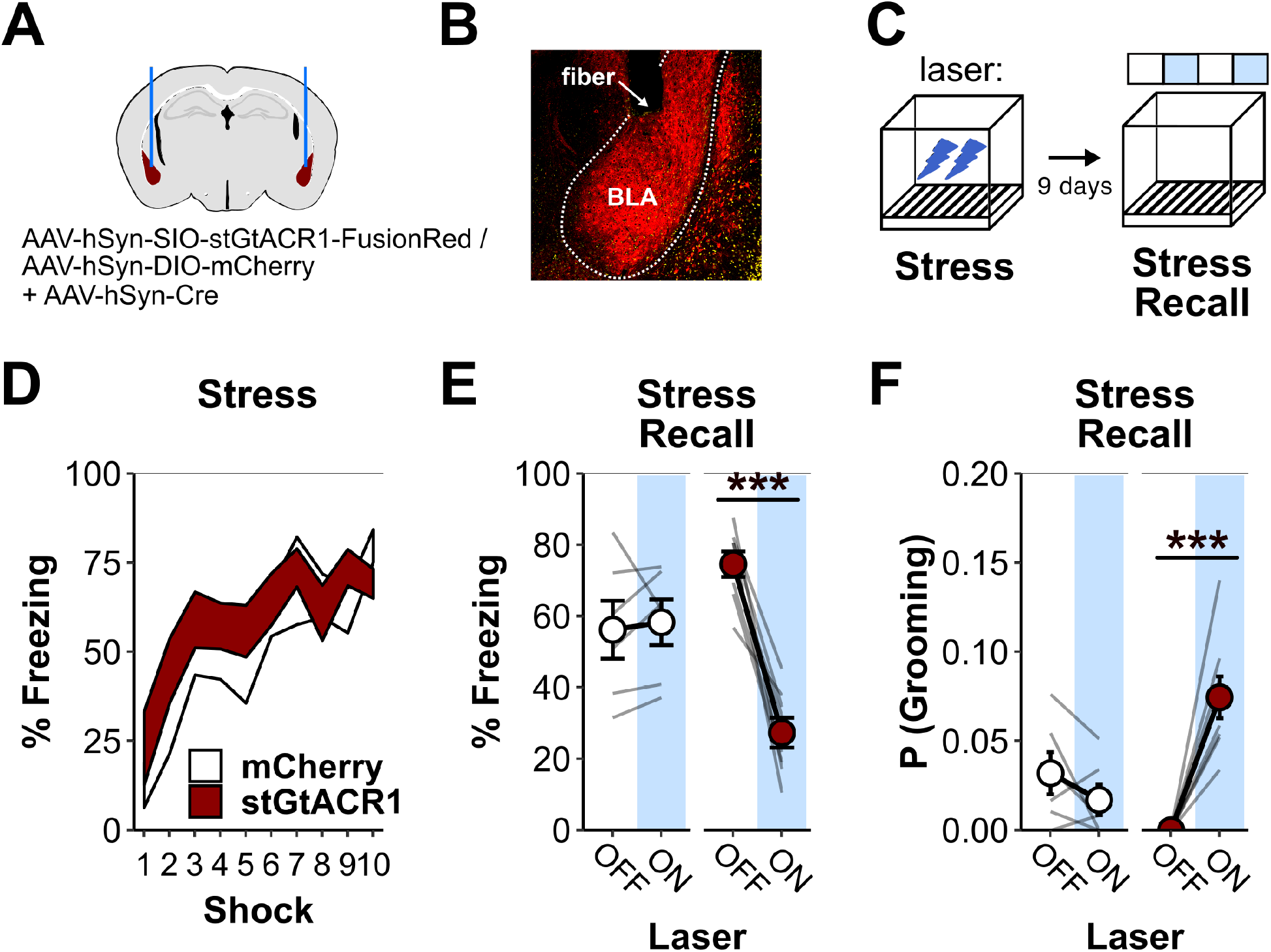
Optogenetic inhibition of stress behaviors restores grooming. A) The inhibitory opsin (stGtACR1), or a control fluorophore (mCherry), were virally expressed in the BLA and bilateral optic cannulas were implanted just above. mCherry: N=6 (all male); stGtACR1: N=8 (all male). B) Example image showing stGtACR1 expression in the BLA and the fiber cannula tip. C) Mice were given a strong footshock stressor. They were subsequently returned to the environment for a recall session, during which the laser was alternated OFF and ON. D) Mice expressing stGtACR1 and control mice increased freezing across the stressor session at a similar rate. Effects of Shock: F_9,108_=13.6, p<0.001; Virus: F_1,12_=0.7, p=0.43; Shock × Virus: F_9,108_=0.7, p=0.71 E) Laser light selectively decreased freezing in mice expressing stGtACR1. Effect of laser for each group: mCherry: F_1,5_=0.17, p=0.7; stGtACR1: F_1,7_=62.6, p<0.001. F) Laser light selectively increased grooming in mice expressing stGtACR1. Effect of laser for each group: mCherry: F_1,5_=2, p=0.22; stGtACR1: F_1,7_=40.3, p<0.001.

### Anxiogenic contexts suppress grooming

Stress-evoked behavior has been theorized to occur along a continuum, with different behaviors emerging as threat levels increase [31]. Moreover, both footshock and footshock-predictive environments may lie on one end of this continuum. To explore the possibility that even lower stress levels might increase grooming, we assessed grooming in the Light-Dark test, a test that is thought to elicit low-level threat or anxiogenic states [32,33]. Additionally, evidence indicates that this form of threat-associated behavior is dependent upon distinct neuronal structures from those necessary for freezing to stress-associated contexts [34,35]. Accordingly, we reasoned that the Light-Dark test may influence grooming differently than footshock and footshock-associated stimuli. As a comparison, we assessed animals’ behavior in a version of the test in which both chambers were dark (Dark-Dark test).

Mice were first habituated to a dark compartment (Side B), which could later be connected to a different environment via a small door in one wall. Then, across two days, they were tested for their preference between the familiar dark environment and a novel light environment (Light-Dark test), and their preference between the familiar dark environment and a novel dark environment (Dark-Dark test) (Fig 3A). In the Light-Dark test, mice spent less time than expected by chance in the light side (Side A, Fig 3B), demonstrating reduced preference for this side. By comparison, in the Dark-Dark test, mice spent a roughly equivalent amount of time in the two sides (Fig 3B). Examining the proportion of time spent grooming on each side, in the Light-Dark test, mice spent significantly less time grooming on the light side than the dark side (Fig 3C). Conversely, grooming did not differ between the two sides of the Dark-Dark test (Fig 3C). Thus, even environments with low threat potential appear to suppress grooming.

**Figure 3.**
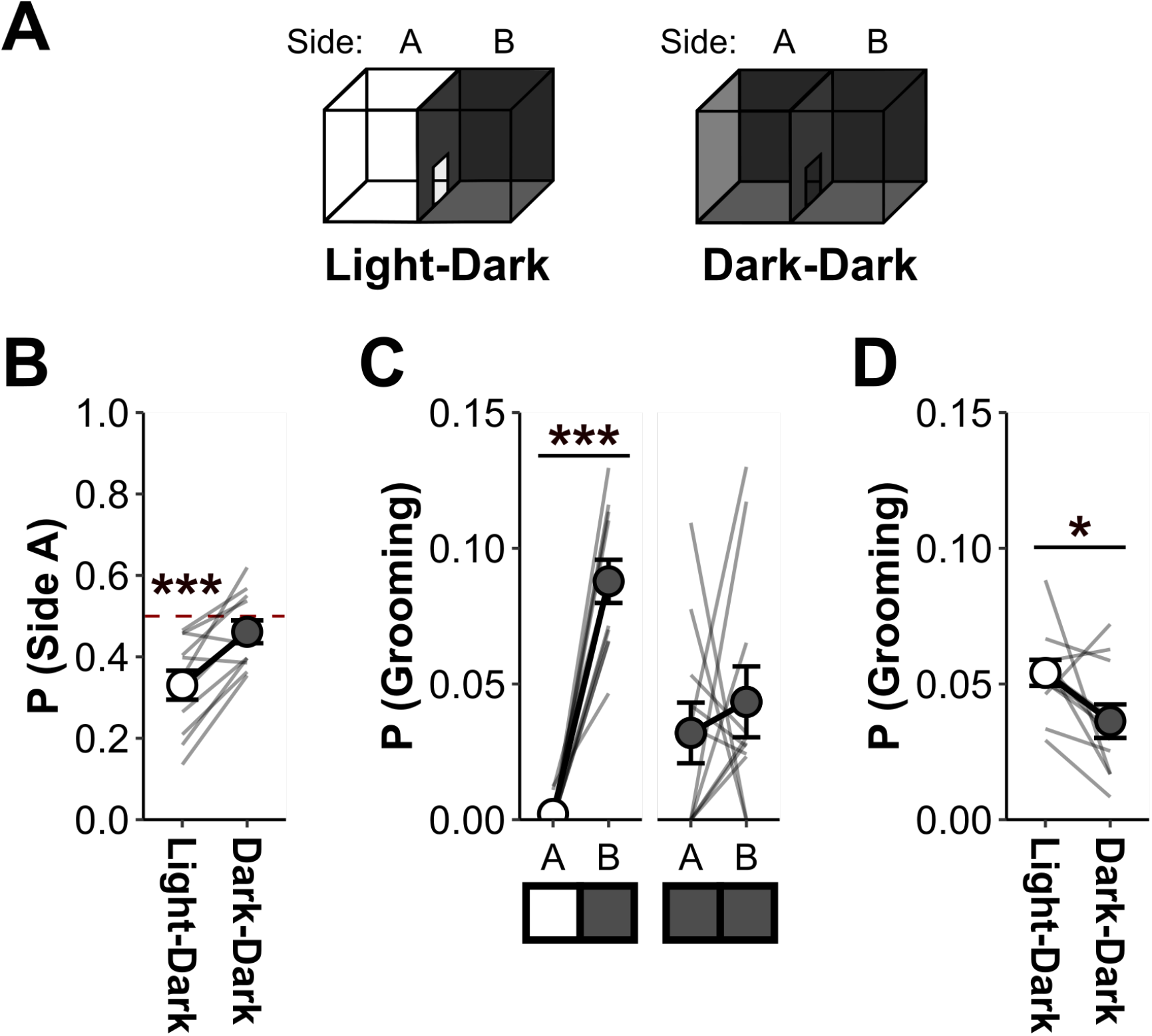
Anxiogenic contexts suppress grooming. A) Mice were assessed in the Light-Dark test as well as an altered version in which both sides were dark (Dark-Dark test). N=11 (6 males, 5 females). B) In the Light-Dark test, mice avoided the light. Conversely, in the Dark-Dark test, no side preference was observed. One sample t-test relative to chance for each test condition: Light-Dark: t_10_=4.7, p<0.001; Dark-Dark: t_10_=1.4, p=0.2). C) In the Light-Dark test, grooming was suppressed on the light side. Effect of Side on time spent grooming for each condition: Light-Dark: F_1,10_=0.3, p=0.59; Dark-Dark: F_1,10_=98.5, p<0.001. Effect of Test condition on time in B: F_1,10_=8.4, p=0.02. D) Overall grooming in the Light-Dark test was higher than in the Dark-Dark test. Effect of test condition on overall grooming: F_1,9_=5.3, p=0.047.

Interestingly, grooming on the familiar, dark, side of the Light-Dark test (Side B) was higher than in the familiar, dark, side of the Dark-Dark test (Fig3C). This could reflect an effort by mice to maintain a homeostatic level of grooming across the session. That is, in the Light-Dark test, mice may groom more in the dark to compensate for less time grooming in the light. If true, the total amount of time spent grooming across the Light-Dark and Dark-Dark test sessions would be expected to be equivalent. However, the total time spent grooming was higher in the Light-Dark test than in the Dark-Dark test (Fig 3D). This suggests that grooming is indeed increased by exposure to a low-stress situation, consistent with prior reports, but only manifests when mice return to a safe environment.

### Grooming increases after stress

The preceding sections highlight that grooming is potently reduced by a range of stressful experiences. Additionally, this suppression is dependent upon stressor strength. Moreover, although grooming on the light side of the Light-Dark test was suppressed, grooming on the dark side increased. This suggested that grooming may increase *after* exposure to a stressful environment, when mice return to a safe environment. This is in line with prior reports showing that grooming increases after a range of stressors, when animals are placed in a distinct environment, though these studies did not directly assess the impact of context on stress-evoked grooming [8,9,11,14,15,23].

To directly test this possibility, we exposed mice to the same footshock stressor that we previously demonstrated reduces grooming behavior. However, this time we also recorded grooming in the homecage immediately after the shock session (Fig 4A). During the stressor session, we again found that Low Stress and High Stress groups showed a suppression of grooming (Fig 4C), along with a complementary increase in freezing levels (Fig 4B). However, when mice were returned to their homecages, the opposite was observed – grooming increased in proportion to the level of stress experienced (Fig 4D).

**Figure 4.**
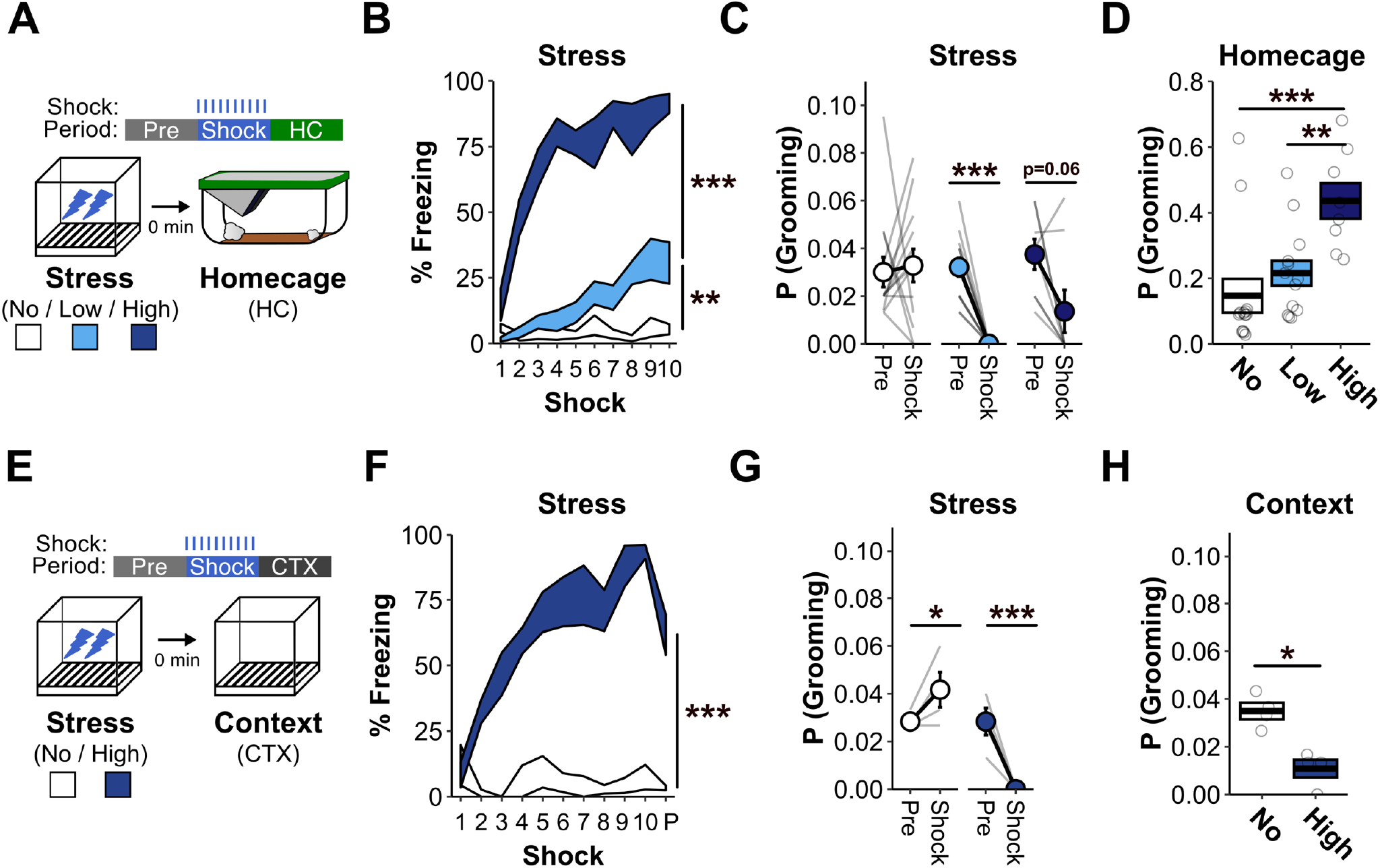
Grooming increases after stress. A) Mice were exposed to No, Low, or High Stress (no shocks, 10 weak shocks, or 10 strong shocks). They were then returned to their homecages and recorded for 10 minutes. No Stress: N=13 (6 females); Low Stress: N=13 (6 females); High Stress: N=8 (4 females). B) Post-shock freezing increased in stressed mice relative to non-stress mice. No Stress vs Low Stress: F_1,22_=10.9, p<0.001. Low Stress vs High Stress: F_1,17_=103.5, p<0.001. C) Grooming was suppressed by both Low Stress and High Stress. Effect of Time for each Group: No Stress: F_1,11_=0, p=0.87; Low Stress: F_1,11_=64.6, p<0.001; High Stress: F_1,6_=5.1, p=0.06. D) Stressed mice displayed heightened grooming levels in the homecage after the stressor. No Stress vs Low Stress: F_1,22_=0.9, p=0.36; No Stress vs High Stress: F_1,17_=16.2, p<0.001; Low Stress vs High Stress: F_1,17_=12.6, p<0.01. E) Mice were exposed to No Stress or High Stress. They were then left in the stressor context for 10 minutes. No Stress: N=4 (2 females); High Stress: N=4 (2 females) F) Post-shock freezing increased in High Stress mice relative to No Stress mice. P corresponds to freezing in the post-shock period. Effect of Group: F_1,4_=98.4, p<0.001. G) Grooming was suppressed during the stressor. Effect of time for each group: No Stress: F_1,2_=32, p=0.03; High Stress: F_1,2_=57.8, p=0.02. H) Grooming continued to be suppressed during the 10 minutes after the stressor. Effect of Group: F_1,4_=16.8, p=0.02.

It could be that the mere passage of time after stress is sufficient to promote the increase in grooming observed, rather than the return to the mice’ homecage. To test this possibility, we performed a similar experiment as the previous one, but instead of returning the mice to their homecages after shock, we left them in the stressor context for an equivalent amount of time (Fig 4E). For this experiment, only No Stress and High Stress groups were used because they showed the greatest difference in grooming in our prior experiments. Here, we found that in the period after shock, High Stress mice continued to not groom (Fig 4G). Therefore, it seems that a context shift is necessary for stress to result in an increase in grooming.

## DISCUSSION

Above, we demonstrate that rodent self-grooming behavior fluctuates in a predictable and bidirectional manner in response to stress. Upon stressor presentation, grooming is immediately suppressed, and this suppression mirrors the intensity of the stressor. Subsequently, when mice return to a relatively safe location, grooming increases to above baseline levels. These findings can be integrated with ethological accounts of rodent defensive behavior to explain past discrepancies in the literature and to foster new predictions. Moreover, these findings have implications for interpreting changes in rodent grooming when modeling aspects of neuropsychiatric function.

Rodents naturally engage in a range of defensive behaviors that serve to protect them from environmental threats or stressors. These include the classic responses of fighting, fleeing, and freezing [31]. The specific defensive behavior elicited in response to a threat has been theorized to depend upon the perceived proximity of that threat and the behavior’s evolutionary advantage to counter that threat level [31,32,36]. For example, when a rodent initially spots a predator lurking in its vicinity, it begins to freeze in order to prevent detection by the predator [37]. This is likely the optimal response, since the rodent stands little chance of either outrunning or fighting off many predators (e.g., a hawk). However, if the rodent is detected and the predator is in close range, the rodent then engages in fight-or-flight behaviors as a last-ditch effort to survive [37]. Now consider grooming behaviors. In the presence of a predator, the initiation of grooming appears to be in clear opposition to the animal’s survival. First, the motion of the rodent would help the predator detect it. Second, the rodent directing attention towards its body and away from the predator provides a prime opportunity for the predator to pounce. Third, grooming and fighting/fleeing are incompatible behaviors. Our finding that the initial response to stress is a suppression of grooming agrees with this ethological view.

Although theories of defensive behavior are more commonly discussed with respect to how behavior changes with escalating threat levels, they also recognize the existence of recuperative behaviors (e.g., wound healing) that help restore the animal after an encounter with a threat [37]. Our findings that grooming increased after stress suggest that grooming may serve a similar restorative role. For example, grooming could prevent the accumulation of bacteria in any wounds obtained during escape from the stressor, or soothe the animal, lessening any negative affective state that accompanies a threatening encounter. Although definitive proof for a restorative function of grooming after stress is still needed, it is interesting that mice will come to prefer a location that has been paired with stimulation of neurons that promote grooming [9], suggesting that grooming has hedonic properties.

This simple account of the relationship between stress and grooming explains the data presented here as well as prior discrepancies in the literature. Although many reports suggest that stress increases grooming, in most of these cases, grooming was measured in an environment distinct from where the stressor occurred [8,9,11,14,15,23]. Accordingly, post-stress behaviors are anticipated. By comparison, when reductions in grooming have been noted in response to stress, grooming was measured before the animal was removed from the stressor environment [19–22]. Therefore, when considering whether grooming is expected to increase or decrease following a stressor, the context in which grooming is measured should be considered.

An alternative account of the bidirectional relationship between stress and grooming is that low stress levels increase grooming, whereas high stress levels decrease grooming [30]. Our results are at odds with this view. We found that grooming was suppressed across a range of stressor intensities. That said, our account may also reconcile why low-stress situations are occasionally be associated with increased grooming. One feature of low-stress situations is the availability of relatively safe locations that the animal can escape to. These safe locations could then provide a place for the animal to engage in post-stress grooming. For example, although we found that grooming on the light side of the Light-Dark test was suppressed (i.e., the stress-associated side), we found that grooming on the dark side was greatly increased (i.e., the safe/preferred side). Indeed, overall levels of grooming in the Light-Dark test were higher than in a similar version of the task with two dark sides. Had we only looked at overall grooming, the suppression of grooming by the light side would have been masked.

Grooming has frequently been used as a phenotype of interest in the study of mental health [2] and our results are of relevance here as well. First and foremost, counter to the prevailing view that grooming is positively correlated with stress and consequently a sign of stress, it is clear that the immediate response to stress is a decrease in grooming. Therefore, in the presence of a stressor, heightened stress levels are denoted by decreased grooming. Furthermore, although stress eventually leads to an increase in grooming, this increase depends upon the removal of the stressor. On this basis, the case could be made that grooming reflects the absence of stress. Alternatively, as we have suggested, the increase in grooming following stress may serve to reduce internal stress levels that remain after the stressor disappears. In either case, the removal of the stressor and the concomitant shift to heightened grooming likely corresponds with a broad change in the animal’s internal motivational state. This shift may be from one of defense to one of recuperation. Moreover, each of these states could be relevant to our understanding of mental health processes.

It is noteworthy that rodent self-grooming has been shown to frequently occur in a stereotyped rostral-to-caudal progression [1,11,15]. Additionally, others have shown that stress is able to alter the temporal organization of grooming. Here, we examined overall levels of grooming, in part because in many of our experiments, the experience of stress led to the complete cessation of grooming, making it impossible to examine the temporal pattern of grooming. That said, in the future, we hope to explore the pattern of grooming during the post-stress period. Given the availability of deep-learning tools that can help tease apart nuanced differences in behavior [38–40], we hope to find signatures of post-stress grooming that can distinguish it from grooming that occurs for a host of other reasons. Such distinctions may reveal a behavioral signature of recuperation.

Our findings also suggest that opposing neural circuits might support what might be called defensive versus recuperative behaviors. The amygdala has long been known to support the learning and expression of defensive responses, including freezing and flight [32,41–45]. Here we found that inhibition of the amygdala not only reduces freezing when mice are placed in an environment previously paired with threat but also results in a dramatic increase in grooming. Accordingly, the neural circuits that support freezing seem to be antagonistic to those that support grooming. This has important implications for studying stress-related disorders. If freezing and stress-induced grooming are controlled by different circuits, then perhaps the human phenotypes they are proposed to model are as well.

In closing, a large amount of effort has been focused on understanding how animals respond to stress at both behavioral and biological levels. By comparison, much less attention has been paid to how animals return themselves to homeostasis once a stressor has abated. In part, this is likely because clear behavioral markers of post-stress recuperation have been lacking, in contrast to the many well-defined behaviors evoked in initial response to stress. Our findings suggest that post-stress grooming could provide such a marker, opening a window into the vital process of how we recuperate.

## AUTHOR CONTRIBUTIONS

ANM and ZTP conceived the overarching research goals, designed the experiments, and oversaw the experiments. ANM, AKP, NY, and ZTP performed experiments. ANM and ZTP analyzed the experimental data and prepared the initial manuscript. ANM, AKP, NY, DJC, and ZTP contributed to interpretation of the results and edited the manuscript. ZTP designed code for analysis of data. ZTP and DJC secured funding.

